# Skeletal myotubes expressing ALS mutant SOD1 induce pathogenic changes, impair mitochondrial axonal transport, and trigger motoneuron death

**DOI:** 10.1101/2024.05.24.595817

**Authors:** Pablo Martínez, Mónica Silva, Sebastián Abarzúa, María Florencia Tevy, Enrique Jaimovich, Martha Constantine-Paton, Fernando J Bustos, Brigitte van Zundert

**Affiliations:** Institute of Biomedical Sciences (ICB), Faculty of Medicine & Faculty of Life Sciences, Universidad Andres Bello, Santiago, Chile; Center for Exercise, Metabolism and Cancer, Facultad de Medicina, Instituto de Ciencias Biomédicas, Universidad de Chile, Santiago, Chile; INTA, Universidad de Chile, Santiago, Chile; McGovern Institute for Brain Research, Department of Brain and Cognitive Sciences, Massachusetts Institute of Technology, Cambridge, MA, USA; Millennium Nucleus of Neuroepigenetics and Plasticity (EpiNeuro), Santiago, Chile; Department of Neurology, University of Massachusetts Chan Medical School (UMMS), Worcester, MA, USA

**Keywords:** ALS, myotubes, muscle, motoneuron, axonopathy, mitochondria, pathology

## Abstract

Amyotrophic lateral sclerosis (ALS) is a fatal neurodegenerative disease characterized by the loss of motoneurons (MNs), and despite progress, there is no effective treatment. A large body of evidence shows that astrocytes expressing ALS-linked mutant proteins cause non-cell autonomous toxicity of MNs. Although MNs innervate muscle fibers and ALS is characterized by the early disruption of the neuromuscular junction (NMJ) and axon degeneration, there are controversies about whether muscle contributes to non-cell-autonomous toxicity to MNs. In this study, we generated primary skeletal myotubes from myoblasts derived from ALS mice expressing human mutant SOD1^G93A^ (termed hereafter mutSOD1). Characterization revealed that mutSOD1 skeletal myotubes display intrinsic phenotypic and functional differences compared to control myotubes generated from non-transgenic (NTg) littermates. Next, we analyzed whether ALS myotubes exert non-cell-autonomous toxicity to MNs. We report that conditioned media from mutSOD1 myotubes (mutSOD1-MCM), but not from control myotubes (NTg-MCM), induced robust death of primary MNs in mixed spinal cord cultures and compartmentalized microfluidic chambers. Our study further revealed that applying mutSOD1-MCM to the MN axonal side in microfluidic devices rapidly reduces mitochondrial axonal transport while increasing Ca2+ transients and reactive oxygen species (i.e., H_2_O_2_). These results indicate that soluble factor(s) released by mutSOD1 myotubes cause MN axonopathy that leads to lethal pathogenic changes.

## 1 INTRODUCTION

Amyotrophic lateral sclerosis (ALS) is a progressive and fatal neurodegenerative disease characterized by the loss of upper and lower motoneurons (MNs), muscle wasting, and paralysis (Al-Chalabi & Hardiman, 2013; Peters et al., 2015; Taylor et al., 2016). Some cases of ALS arise in association with frontotemporal dementia (FTD) (Ling et al., 2013). Sporadic ALS (sALS) cases are responsible for most of the cases (90%), while the remaining 10% have a familial history of ALS (fALS), characterized by genetic inheritance (Renton et al., 2014). Approximately 20% of fALS corresponds to mutations in the superoxide dismutase 1 (mutSOD1) gene, where more than 160 mutations have now been identified in its sequence (http://alsod.iop.kcl.ac.uk/). The discovery of SOD1 mutations in fALS patients (Rosen et al., 1993) led to the generation of the first transgenic ALS mouse model expressing SOD1^G93A^ (Gurney et al., 1994). As of today, the high copy number SOD1^G93A^ transgenic mouse model is still a cornerstone of ALS research because this model closely recapitulates the human clinical and histopathological symptoms of ALS and exhibits a stable and well-established disease progression, enabling preclinical testing of novel genes and pharmacological therapies (Zundert & Brown, 2017).

Based on studies *in vitro* with cultures and *in vivo* with mouse models, it is widely accepted that MN degeneration in ALS occurs through non-cell-autonomous mechanisms, involving interactions between various local cell types such as astrocytes, microglia, and oligodendrocytes (Ilieva et al., 2009; Maimon & Perlson, 2019). Particularly, there is compelling evidence that ALS astrocytes cause non-cell-autonomous toxicity to MNs (Dittlau & Bosch, 2023; Garcés et al., 2024; Harten et al., 2021). For example, in *in vivo* studies, the survival of mutSOD1 mice is significantly extended when ALS MNs are surrounded by wild-type (WT) non-neuronal cells (Clement et al., 2003), particularly astrocytes (Lepore et al., 2008). Selective deletion of the mutSOD1 genes from astrocytes markedly prolongs the survival of the mutSOD1 mice by delaying disease onset and/or progression (L. Wang et al., 2011; Yamanaka et al., 2008). Conversely, studies in rodents, either by selectively expressing ALS-linked mutant genes or by transplanting glial progenitors, suggest that fALS astrocytes (Papadeas et al., 2011; Tong et al., 2013), and sALS astrocytes (Qian et al., 2017) can induce certain aspects of MN degeneration, locomotor deficits, and decrease survival. Non-cell autonomous toxicity to MNs induced by cultured astrocytes harboring diverse ALS-causing gene mutations, including mutations in SOD1, TDP43, and C9ORF72, has also been extensively documented (Dittlau & Bosch, 2023; Garcés et al., 2024; Harten et al., 2021). Our recent study further reveals that excessive inorganic polyphosphate (polyP) released by ALS astrocytes triggers MN death and increases neuronal excitability and Ca^2+^ transients (Arredondo et al., 2022; Garcés et al., 2024; Rojas et al., 2023). Additionally, other studies have also shown that astrocyte-mediated MN degeneration is accompanied by several other pathogenic events, including oxidative stress, induction of a cell death signaling (i.e., phosphorylation of c-Abl), and impaired mitochondrial transport (Birger et al., 2019; Dittlau & Bosch, 2023; Fritz et al., 2013; Haidet-Phillips et al., 2011; Harten et al., 2021; Rojas et al., 2014, 2015).

In addition to astrocytes, studies in patients and animal models indicate that cells outside the central nervous system (CNS) are also affected in ALS, including lymphocytes (Cova et al., 2006), fibroblasts (Aguirre et al., 1998), and skeletal muscle (Cappello & Francolini, 2017; Dobrowolny et al., 2008; Frey et al., 2000; W. Guo et al., 2020; Hegedus et al., 2007; Jensen et al., 2016; Wong & Martin, 2010). Regarding the latter, while skeletal muscle cells innervate muscle fibers and ALS is characterized by the early disruption of the neuromuscular junction (NMJ) and axon degeneration (Cappello & Francolini, 2017; Fischer et al., 2004; W. Guo et al., 2020; Moloney et al., 2014), there are controversies whether muscle contributes to MN degeneration. While reducing SOD1 levels directly in the muscles of mutSOD1 transgenic mice did not affect the onset of the disease or survival (T. M. Miller et al., 2006) muscle-restricted expression of mutSOD1 led to alterations associated with ALS pathogenesis (Dobrowolny et al., 2009) and a classic ALS mouse phenotype (Dobrowolny et al., 2009; Maimon et al., 2018; Wong & Martin, 2010). *In vitro*, studies also report controversial results with data favoring (Maimon et al., 2018) or contrasting (Nagai et al., 2007) evidence for showing non-cell-autonomous toxic actions of mutSOD1 myotubes to healthy MNs.

Here, we demonstrate that myotube-conditioned media (MCM) derived from primary myotubes harboring mutated human SOD1^G93A^ (mutSOD1-MCM) induce MN death and trigger the accumulation of ROS and phosphorylated c-Abl. Furthermore, we found that distal application of mutSOD1-MCM increases intracellular Ca^2+^ events, ROS production, and death of primary wild-type MNs using microfluidic devices. We also found that MCM-mutSOD1 produces a functional deficit of axonal mitochondrial transport, both retrograde and anterograde. Our findings provide compelling evidence that myotubes contribute to MN degeneration in SOD1-linked ALS.

## 2 MATERIALS AND METHODS

### 2.1 Mice handling

All mice used were handled according to the guidelines for the handling and care of experimentation established by the NIH (NIH, Maryland, USA) and following the protocol approved by the bioethics committee of Andres Bello University (approval certificate 014/2017). We used hemizygous transgenics mice harboring the human mutation SOD1^G93A^ (High number of copies; B6SJL) obtained from Laboratories Jackson (Cat. No. 0022726, Bar Harbor, ME, USA). Non-transgenic littermates were used as controls. The presence of the transgene was identified by end-point PCR (Fritz et al., 2013).

### 2.2 Myoblasts and myotubes cultures

Cultures of myoblasts and myotubes were made from postnatal days 1-4 (P1-4) pups as described previously (Valdés et al., 2007). Briefly, pups were euthanized by decapitation and left in PBS 1X (Hyclone, No.SH30538.03). Muscle tissue was obtained from lower limbs and incubated for 15 min at 37℃ in a collagenase type-2 (1.5 mg/mL; Gibco, No.17101-015), dissolved in PBS 1X and filtered with 0.22 μm filters. Then, the tissue was mechanically disintegrated and incubated for 15 min at 37℃ with the collagenase solution. Subsequently, it was mechanically disintegrated for a second time, and 5 ml of F-10 medium was added (Sigma, No. 6635-1L). The solution was filtered through 40 μm filters (Falcon, No. 352340) and centrifuged at 1100 RPM for 7 min. The supernatant was removed, and the pellet was resuspended in myoblast growing media (F-10 supplemented with 20% bovine growth serum (BGS, Hyclone, No. SH30541.03), 1% penicillin/streptomycin (Gibco, No. 15114-122), and five ng/mL final concentration of human fibroblast growth factor (FGF; PeproTech, No. 100-18B-100UG). Cells were plated in 100 mm plates (without any matrix) for 45 min at 37℃ and 5% CO2. After this time, adhered fibroblasts were observed, and the supernatant containing the myoblasts was removed and plated on a new 100 ml plate treated with Matrigel (Sigma, No. E1270). The culture medium was replaced every two days. When the myoblasts reached 70% confluency (3-5 DIV), growing media was replaced by differentiation media that consisted of DMEM (Gibco, No. 11885-084) supplemented with 10% fetal bovine serum (FBS; Gibco, No. 16000-044), 4% horse serum (Gibco, No. 16050), 1% L-glutamine (Gibco, No. 25030) and 1% penicillin/streptomycin. The fusion of myoblasts was observed from the second day after the medium change. The differentiation medium was replaced every four days until the complete fusion and differentiation to myotubes (∼10 DIV after medium change), which showed spontaneous contractions under the light microscope.

### 2.3 Spontaneous contraction assay

Mature myotube cultures were visualized using an epifluorescence microscope (Nikon Eclipse Ti-U; objective 20X) 10 DIV after differentiation. Spontaneous contractions were observed and quantified manually and normalized to 1 min.

### 2.4 Preparation of myotube-conditioned media

Conditioned media was prepared from primary cultures of myotubes. At 80-90% of confluence (8-10 DIV after differentiation induction) culture medium was replaced by ventral spinal cord neurons growth media (Fritz et al., 2013), containing 70% MEM (Life technologies 11090-073), 25% Neurobasal media (Life technologies 21103-049), 1% N2 supplement (Life technologies 17502-048), 1% L-glutamine (Life technologies 25030-081), 1% penicillin-streptomycin (Life technologies 15070-063), 2% horse serum (Hyclone SH30074.03) and 1 mM pyruvate (Sigma). The media was left for seven days, supplemented with D(+)Glucose (Sigma, No.G7021), and filtered with 0.22 μm filters. The myotube conditioning media (MCM) was stored at -80℃ for six months.

### 2.5 Primary ventral spinal cord cultures

Sprague Dawley rats were cared for and handled according to the guideline practices of managing and caring for experimental animals established by the NIH (NIH, Maryland, USA) and following the approval of the Ethics Committee of Andres Bello University. Primary ventral spinal cord cultures were prepared from embryonic day 18 (E18) rats as previously described by our laboratory (Fritz et al., 2013; Sepulveda et al., 2010). Briefly, pregnant wild-type rats were euthanized using a CO2 chamber, and E18 embryos were removed and rapidly decapitated. Tissues were placed in a cold PBS1X solution supplemented with 1% penicillin/streptomycin. The dorsal portion of the cord was removed using a sterile scalpel. The ventral spinal cords were mechanically dissociated and incubated for 20 min at 37℃ in prewarmed 1X PBS supplemented with 0.25% trypsin (Life Technologies 15090-046). After incubation, the cells were transferred to 15 ml tubes containing feeding medium: Minimum Essential Media (MEM; Life Technologies, 11095-072) supplemented with 10% Horse serum (Hyclone, SH30074.03), 1% L-glutamine (Life Technologies, 25030-081), 4 mg/mL DNAase (Roche, 04716728001). The cells were resuspended by mechanical agitation through Pasteur pipettes flamed with decreasing diameters. Cells were counted and seeded (400,000 cells/mL for survival assays) on poly-L-lysine treated glasses (MW 350 kDa, Sigma Chemical, St. Louis, MO). The feeding medium was replaced to growth media: 70% MEM (Life technologies 11090-073), 25% Neurobasal media (Life technologies 21103-049), 1% N2 supplement (Life technologies 17502-048), 1% L-glutamine (Life technologies 25030-081), 1% penicillin-streptomycin (Life technologies 15070-063), 2% horse serum (Hyclone SH30074.03) and 1 mM pyruvate (Sigma S8636). The cultures were supplemented with 45 μg/ml of E18 chicken leg extract kept at 37 ℃ and 5% CO2, with medium replaced every 3 days.

### 2.6 Immunofluorescence

For myoblast and myotube cultures characterization, MN survival, and c-Abl phosphorylation, immunofluorescence assays were performed on 4% PFA (20 min) fixed cultures, followed by three washes with PBS 1X. Then, the cells were permeabilized with Triton X-100 at 0.05% v/v in PBS 1X for 30 min and washed three times with PBS 1X. Cells were blocked for 30 min with goat serum (Life Technologies, No. 50062Z). Cells were incubated with the different primary antibodies overnight at 4℃. Primary antibodies used: Pax7 (DSHB, AB_528428, 1:100), SMI32 (Abcam, Ab187374, 1:500), MAP2 (Merck Millipore, Mab378, 1:250), MyoD (DSHB, D7F2-s, 1:500), MHC (Novus Biologicals, NB300-284, 1:1000), Myogenin (Abcam, Ab1835, 1:500) and c-Abl Tyr-412 (SIGMA, C5240, 1:1000). The next day, cells were washed three times with PBS 1X for 5 min each wash and incubated with the secondary antibodies conjugated to Alexa 488, Alexa 555, or Alexa 633 for 2.5 h at room temperature. Secondary antibodies used: goat anti-rabbit Alexa Fluor 488 (Life Technologies, A11008, 1:500), goat anti-mouse Alexa Fluor 488 (Life Technologies, A10667, 1:500), goat anti-rabbit Alexa Fluor 555 (Life Technologies, A21428, 1:500), goat anti-mouse Alexa Fluor 555 (Life Technologies, A21422, 1:500), goat anti-rabbit Alexa Fluor 633 (Life Technologies, A21070, 1:500), goat anti-mouse Alexa Fluor 633 (Life Technologies, A21050, 1:500). In parallel to the secondary antibodies’ incubation, the cells were incubated with DAPI (Sigma, No. D9542). Cells were then washed three times for 5 min in PBS 1X and mounted using Fluoromont G fluorescence mounting medium (EMS, No. 17984-25). For all immunofluorescence analysis, we used an epifluorescence microscope (Nikon Eclipse Ti-U; objective 20X).

### 2.7 MN survival assay

Survival of MNs was measured as previously described by our laboratory (Fritz et al., 2013; Rojas et al., 2014). Briefly, spinal cord cultures were fixed at 7 DIV, and immunofluorescence was performed using the above protocol. We used a specific antibody for MAP2 to detect all neurons in the cultures (including interneurons and MNs) and a specific antibody for SMI32 to identify only MNs (Arredondo et al., 2022; Fritz et al., 2013; Mishra et al., 2020; Nagai et al., 2007; Sepulveda et al., 2010). Previously, we have described that primary spinal cord cultures contain 8-10% of MNs at 12 DIV (Sepulveda et al., 2010). Fluorescent staining was visualized by epifluorescence microscopy (Nikon Eclipse Ti-U; objective 20X). The fields with neurons were randomly chosen, and the number of MAP2+ and SMI32+ neurons was counted from all the acquired images. Per condition, ≥ ten randomly selected fields (≥200 cells) were analyzed to calculate the percentage of SMI32+ MNs for the total number of MAP2+ cells. The ratio between SMI32^+^/MAP2^+^-cells and SMI32^-^/MAP2^+^ neurons indicate the percentage of MN survival compared to the control media. Each condition was replicated in 3-4 independent cultures.

### 2.8 Reactive oxygen species (ROS) production assay

Intracellular ROS levels were measured as previously described by our laboratory (Rojas et al., 2015). Briefly, a stock of 5 mM of the CM-H2DCF-DA probe (Invitrogen, Cat. No.C6827) was prepared fresh in DMSO and then diluted in the culture medium to a final concentration of 1 μM. Cells were washed with PBS 1X to remove the different MCMs for 90 min after applying the different conditioned medium and the CM-H2DCF-DA probe for 30 min at 37 ℃ in the dark. To facilitate the incorporation of the probe into cells, 0.004% pluronic acid F-127 (Invitrogen, Cat. No. P-3000MP) was added. After the incubation, the probe CMH2DCF-DA dissolved in the culture media was removed, and the cells were washed twice with PBS 1X to apply the culture medium to the spinal cord neurons. Cultures were also incubated with H_2_O_2_ (200 μM for 20 min) as a positive control to normalize the number of positive DCF cells after the insult with MCMs. Imaging was made using an epifluorescence microscope (Nikon Eclipse Ti-U; objective 20X) and excitation and emission wave λex/λem = 492–495/517–527 nm. At least three fields were taken for each condition, and at least ten cells per field were used for the quantification. The analysis of images was done using ImageJ software (NIH, Bethesda, MD, USA).

### 2.9 phosphorylated c-Abl (c-Abl-P) immunofluorescence labeling

c-Abl phosphorylation in MNs was determined by immunofluorescence labeling as previously described (Rojas et al., 2014). Briefly, primary spinal cord cultures were exposed to the different MCMs, fixed at 7 DIV with 4% PFA, and incubated with antibodies against SMI32. To detect phosphorylated c-Abl, a mouse monoclonal antibody that recognizes phosphorylation of Tyr-412 was used and visualized with the appropriate Alexa fluorescent secondary antibody (see section 2.6 immunofluorescence). For c-Abl-P quantification in cultures, cultures were imaged using a 20X objective. The fluorescence intensity was quantified in SMI32^+^ MNs using ImageJ software (NIH, Bethesda, MD, USA). Briefly, the cell body of each SMI32^+^ neuron was marked manually to set a region of interest (ROI), and the mean c-Abl-P fluorescence was quantified. The background was subtracted, choosing a region without cells. The fluorescence corresponding to control cells was normalized to 1.

### 2.10 Microfluidic system

We use microfluidic chambers with microchannels 450 μm long (Xona, No. SND450). The sterile chambers were mounted on glass coverslips previously treated with poly-L-lysine for 30 min at 37℃. Once adhered, chambers were incubated at 80℃ for 1 h to allow the correct adhesion between the chamber and the glass. Chambers were then exposed to UV light for 5 min for sterilization. 100 μl of the medium was applied to each chamber, and MNs were plated (adapted from (Southam et al., 2013)).

### 2.11 MN and myotube co-cultures

Enriched MN cultures for microfluidic experiments were performed as described previously (Milligan & Gifondorwa, 2011) with some modifications. Briefly, P0-P1 wild-type mice pups were euthanized by decapitation, the skin on the back removed, and the spinal cord was taken, opening each vertebra right down the middle. The spinal cord was left in cold 1X PBS and subsequently transferred to the digestion solution: Papain (Sigma, No. P3375), DNAase (Roche, No. 10104159001), MgCl2 (Sigma, No. M2393), 1 ml PBS 1X. Spinal cords were mechanically disaggregated using tweezers, leaving pieces of approximately 0.3 mm, then incubated at 37℃ for 10 min, to be later transferred to a 15 ml tube and washed three times by adding 1X PBS. The spinal cord pieces were placed in 5 ml of preheated MN medium: Neurobasal Medium (Gibco, No. 21103-049), 1% Glutamax (Gibco, No. 35050-061), 2% Supplement B27 (Gibco, No. 17504-04), 5% of horse serum), and DNAase and MgCl_2_. Then, the marrows were disintegrated using glass Pasteur pipettes, previously flamed at the tip, with two different thicknesses, passing each spinal cord through the pipettes from the greater to the smaller hole. To purify the MNs, an Optiprep gradient column was assembled (Milligan & Gifondorwa, 2011). Cells were centrifuged at 1900 rpm for 15 min. After centrifugation, the third layer of the generated gradient, which contains a higher proportion of MNs was removed. The cells were left in a new 15 ml tube with 10 mL of prewarmed MN medium and were centrifuged at 1000 rpm for 10 min. The pellet was carefully resuspended in no more than 200 μl of preheated MN medium, and cells were counted and diluted up to 1×10^6^ cells/ml. Approximately 5×10^4^ cells were plated in the proximal compartment in no more than 50 μl of MN medium. The cells were incubated for 30 min at 37℃ and 5% CO2 for cell adhesion. Subsequently, 200 μl of MN medium was applied to the culture of MN localized in the upper proximal compartment. The culture medium was replaced every 48 hours. At 3 DIV, the first axonal prolongations were observed, and that was the time that the myoblasts were plated in the distal compartment. Like MNs, 5×10^4^ myoblast cells from the protocol described before (see section 2.2 myoblasts and myotubes cultures) were resuspended in 50 μl of myoblast growth medium and plated in the upper distal compartment. The chamber was incubated for 30 min at 37℃ and 5% CO2, and finally, 200 μl of myoblast growth culture medium was applied in the upper distal compartment. The next day, the medium was replaced to differentiate myotubes to induce fusion of the myoblasts. Approximately between 10 and 14 DIV, it was possible to observe the innervation of MN axons to the distal compartment where the myotubes were cultured.

### 2.12 Adeno-associated viral particle production

Based on previous work (Arredondo et al., 2022; Bustos et al., 2017, Bustos et al., 2023) HEK293FT cells were grown on 150 mm plates in DMEM medium (Hyclone SH30081.02) until reaching 80-90% of confluence and supplemented with 10% fetal bovine serum (FBS; Hyclone SH30070.01), 4 mM L-glutamine (Life technologies 25030-081), 100 U/ml penicillin/streptomycin (Life technologies 15070-063) and 1 mM pyruvate (Sigma), at 37 ℃ and 5% CO2. Cells were transfected using polyethyleneimine (PEI), and the plasmids for adeno-associated viruses type 1 and 2 (pAAV1 and pAAV2) were utilized. In addition, we use the pFΔ6 plasmid and the following plasmids of interest: hSyn-COX8-RFP, a plasmid with the synapsin-1 promoter that controls the expression of COX8, exclusive from mitochondria, and fused to the red fluorochrome RFP; hSyn-mRuby2-GCaMP6, a plasmid with the synapsin-1 promoter that controls the expression of the biosensor GCaMP6 and fused to the red fluorochrome mRuby2 (Addgene, # 50942); and hSyn-HyPer, HyPer biosensor 3.0 under CMV promoter (Addgene, #42131). We transfected 10.4 μg of pFΔ6, 4.35 μg of pAAV1, 4.35 μg of pAAV2, 5.2 μg of the plasmid of interest and 880 µl of Optimem (Gibco, No. 31985-070). Plasmids were mixed, and 260 μl of polyethyleneimine reagent (PEI; Sigma, No. P3143) was added. The solution was applied on 80% confluent HEK293T cells, and after 12 h, the medium was replaced by DMEM medium with 1% FBS. At 72 h from the start of the transfection, cells were collected and centrifuged at 3000 RPM for 10 min 4 ℃. The supernatant was discarded, and the pellet was resuspended in 4 ml 1X PBS. Cells in suspension were left at -80℃ for 10 min and then thawed at 37℃ for 10 min. This cycle was repeated three times to achieve thermal lysis of the cells. Finally, the supernatant was removed and filtered using 1.2 μm, 0.45 μm and 0.22 μm filters. The solution containing the viral particles was aliquoted and refrigerated at -80℃.

### 2.13 Mitochondrial velocity assay

Microfluidic co-cultures of MNs and myotubes were transduced at 10-14 DIV in the proximal compartment using AAV1/2 coding for hSyn-COX8-RFP. 4-7 After transduction (14-21 DIV of co-culture), it was possible to observe the expression of RFP with a mitochondrial pattern in MN under epifluorescence microscopy. At this point, the culture medium was replaced by the different MCMs, and 24 h later the registration of the mitochondrial movement was measured. The microfluidic chamber was placed in a CO_2_ chamber (Tokai-Hit) coupled to an epifluorescence microscope (20X objective; Nikon TE-2000). Sequential images of axons that crossed the microchannels in direct contact with myotubes were recorded. The images were acquired every 2 seconds for 5 min to generate a pattern of the mitochondrial movement. The images were processed in the ImageJ software using the Kymograph plugin. The slopes obtained by Kymograph represent mitochondrial velocity (Zahavi et al., 2015).

### 2.14 Analysis of calcium events in MNs

Co-cultures of MNs and myoblasts were generated in the microfluidics chambers as described above. Next, cells in the proximal compartment containing the MNs were infected with AAV1/2 containing hSyn-mRuby2-GCaMP6s, which encodes for the GCaMP6s biosensor to detect intracellular calcium fluctuations (Chen et al., 2013).After 5-7, DIV of transduction, calcium frequency recordings were performed using an epifluorescence microscope (Nikon TE2000e, 20X objective, and Andor Zyla camera 5.5). Images were taken every 50 ms for 1-1.5 min with an excitation wavelength of 480 nm and emission of 510 nm. The soma of an isolated MN was selected to quantify the frequency of calcium events. Images of the isolated soma were analyzed using the ImageJ software and the Z-Profiler tool, which provides an intensity profile during the time. Each intensity peak was counted manually and was divided by 1 min, resulting in a value expressed as events per min.

### 2.15 Analysis of ROS production in MN in the microfluidic system

Co-cultures of MNs and myoblasts were generated in the microfluidics chambers as described above. Next, cells in the proximal compartment containing MNs were infected with AAV1/2 containing the hSyn-HyPer 3.0 plasmid, which encodes for the HyPer biosensor and YFP protein (Bilan et al., 2013). After 5-7 DIV of transduction, ROS production recordings were performed using an epifluorescence microscope, with an excitation wavelength at 480 nm and emission at 510 nm. The acquisition was made every 2 seconds for 10 min. The images were analyzed using the ImageJ software, using the Z-Profiler tool to obtain the intensity profile over time.

### 2.16 Data analysis

Statistical analyses were performed using GraphPad Prism software. Student’s t-test was performed when two populations were examined, while one-way ANOVA followed by the Bonferroni post-hoc was utilized when making multiple (three or more) comparisons. In all figures, data is reported as mean±S.E.M.; *p < 0.05, **p < 0.01, ***p < 0.001 compared to control. For all data, three or more independent experiments were analyzed.

## 3 RESULTS

### 3.1 MutSOD1 myotubes acquire aberrant phenotypic and functional characteristics

We first aimed to determine the phenotypic characteristics of mutSOD1 myoblasts and myotubes primary cultures. Primary myoblast cultures were generated from lower limb skeletal muscles derived from postnatal day (P) 1-4 neonatal ALS transgenic mice carrying human mutant SOD1^G93A^. As controls, myoblasts were isolated from non-transgenic littermates (NTg). Using immunofluorescence staining assays for Pax7 and myosin heavy chain (MHC), we tested the differentiation of myoblasts to myotubes. Pax7 is essential for the normal expansion and differentiation of satellite cells (SCs) into myoblasts in both neonatal and adult myogenesis (Maltzahn et al., 2013), while MHC is a marker for functional myotubes (X. Guo et al., 2020; Torgan & Daniels, 2001). At 3-5 DIV, when NTg and mutSOD1 myoblasts reached 70% confluence, we observed a robust expression of the transcription factor Pax7 (Fig. 1A). Next, differentiation media was applied to induce complete fusion and differentiation to myotubes. As expected, during the differentiation process the expression of Pax7 decreased concomitantly with an increased expression of MHC (Fig. 1A). Both NTg and mutSOD1 cultures displayed the myoblast to myotube transition over ten days. Next, we analyzed the efficiency of the myotube formation by performing a short differentiation assay (8 h) followed by double immunofluorescence assays to determine the expression of both Pax7 and myotube marker myogenin (MyoG) (González et al., 2016). Interestingly, mutSOD1 myotubes showed a significant reduction (∼50%) in myogenin induction (Pax7^-^/MyoG^+^) compared to NTg myotubes, indicating a deaccelerated commitment to myotube differentiation (Fig. 1B-C). To further characterize the myogenic process in the ALS cells, we analyzed the structural and functional traits of NTg and mutSOD1 myotubes. Since myotube cultures show spontaneous contractions (X. Guo et al., 2013; Smolina et al., 2015), we analyzed the contraction capability in our model. We found that mutSOD1 myotubes displayed a significantly higher contraction frequency (∼55%), compared to NTg myotubes (Fig. 1D). Moreover, we determined that mutSOD1 myotubes display a significantly lower width (∼23%), compared to NTg myotubes (Fig. 1E), without evident changes in nuclei number per myotube (Fig. 1F). Together, these results show that mutSOD1 myotubes display intrinsic phenotypic and functional differences compared to NTg muscle cells.

**Figure 1.**
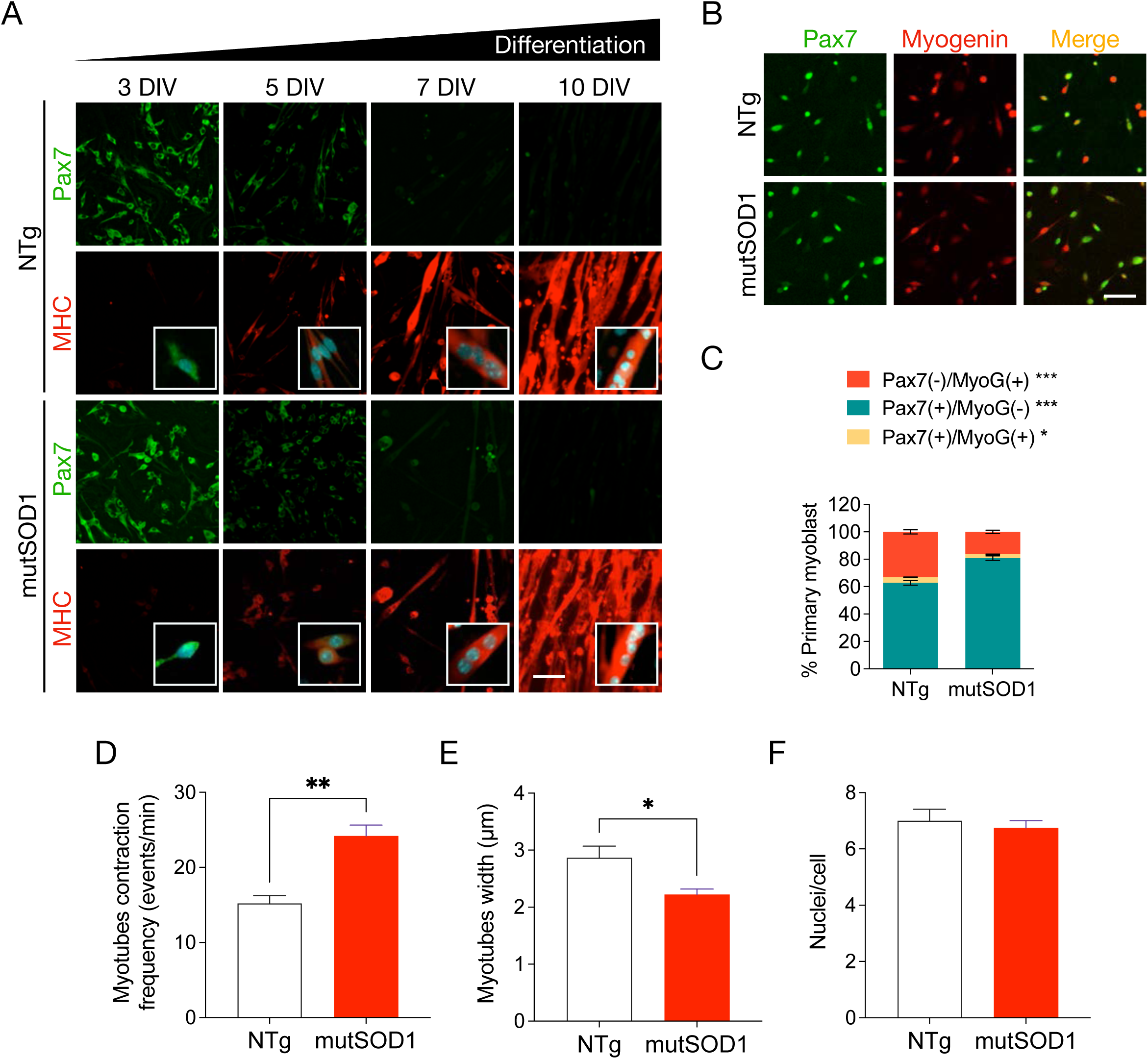
Characterization of primary mutSOD1 myoblast cultures. **A**, Representative images of myogenic markers during myoblast differentiation in primary mutSOD1 and non-transgenic littermate (NTg) myoblast cultures. Myoblasts from P2 mice were maintained in a growth medium up to 70% confluence and then cultured in a differentiation medium to induce myotube formation. Cells were fixed at 3, 5, 7, and 10 DIV and immunostained with antibodies against Pax7 and MHC, and DAPI to detect nuclei (n=3). Scale bar: 100 µm. **B**, Quantification of subpopulations present in primary NTg and mutSOD1 myoblast cultures. Five DIV primary mutSOD1 and NTg myoblasts were induced to differentiate into myotubes for 8 h. Cells were fixed, and immunofluorescence was performed using specific antibodies against Pax7 and myogenin (MyoG). Scale bar: 100 µm. **C**, Pax7, and myogenin-positive (and negative) cells were quantified to obtain the enrichment percentage of each myogenic gene over the total number of cells. The quantification corresponds to 3 independent experiments, analyzed by student t-test (* p <0.05, *** p <0.0005). **D**, Myotube contraction frequency, comparing NTg and mutSOD1 myotubes and quantified as event per min. Data are represented as the mean ± s.e.m., student t-test (** p<0.005). **E**, Myotube width. Comparison made between NTg and mutSOD1 myotubes in 3 independent experiments. Data are represented as the mean ± s.e.m., student t-test (** p<0.005). **F**, Number of nuclei per cell in NTg and mutSOD1 myotubes. The quantification corresponds to 3 independent experiments, analyzed by student t-test. No significant differences were detected.

### 3.2 Exposure to mutSOD1-MCM triggers the death of primary MNs

Next, we aimed to explore the hypothesis that skeletal muscle expressing mutSOD1 causes MN pathology and death by releasing soluble toxic factor(s). We established an *in vitro* culture model (Fig. 2A) in which myotube-conditioned media (MCM) from mutSOD1 myotubes (termed mutSOD1-MCM) was applied at different dilutions to ventral spinal cord cultures at 4 DIV. MCM from NTg astrocytes (termed NTg-ACM) and fresh media to maintain MNs in culture (termed MN medium) were included as controls. At 7 DIV, cultures were fixed and double immunostained for unphosphorylated neurofilament-H (SMI32) and MAP2 to identify MNs (SMI32^+^/MAP2^+^-cells) or interneurons (SMI32^-^/MAP2^+^-cells) (Arredondo et al., 2022; Fritz et al., 2013; Mishra et al., 2020; Nagai et al., 2007; Sepulveda et al., 2010). In agreement with our hypothesis, we found that a 1/4 (25%) and 1/8 (12.5%) dilution of mutSOD1-MCM strongly reduced MN survival (∼40-50%), whereas NTg-MCM did not cause MN death (Fig. 2B-C).

**Figure 2.**
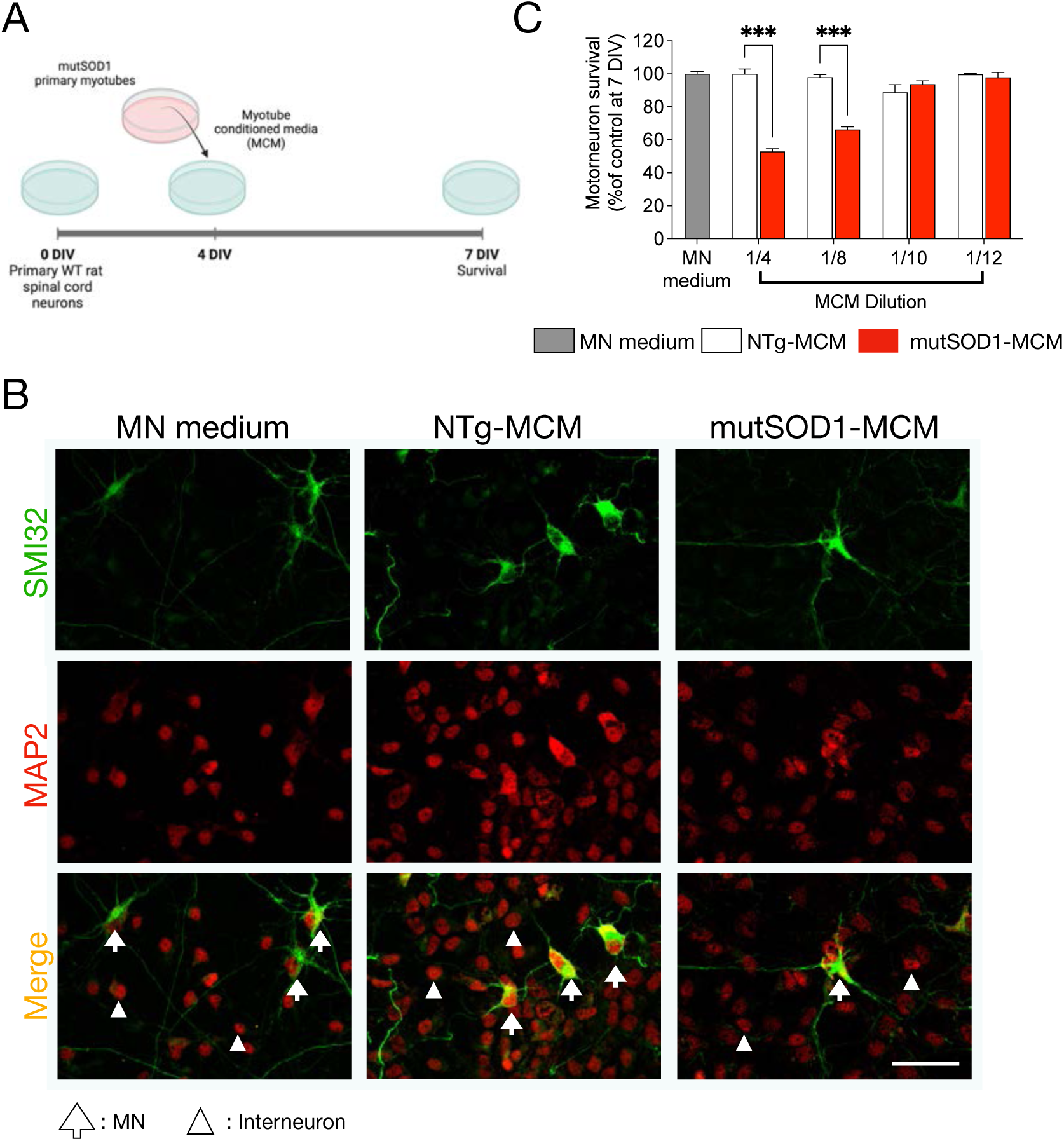
MCM-mutSOD1 contains soluble toxic factor(s) that induce(s) MN death. **A**, Workflow diagram of primary WT (NTg) spinal cord cultures (4 DIV) that were exposed for 3 days to MCM derived from mutSOD1 transgenic mice (MCM-mutSOD1), NTg littermates (NTg-MCM), and culture medium (MN medium). Cells were fixed at 7 DIV, and immunofluorescence assayed cell survival. **B**, Representative images of immunofluorescence against SMI32 (MNs) and MAP2 (all neurons) when exposed to MCM-mutSOD1 (dilution 1/4), NTg-MCM (dilution 1/4), and MN medium. Scale bar: 50 μm. **C**, MN survival graph (SMI32^+^/MAP2^+^ cells as a percentage of all MAP2^+^ neurons) after treatment with MCM-mutSOD1, NTg-MCM, and MN medium for 3 DIV. Values represent the mean ± s.e.m of at least three independent experiments performed in duplicate and analyzed by one-way ANOVA (*** P <0.0005) relative to the NTg-MCM at 7 DIV.

### 3.3 Application of mutSOD1-MCM to spinal cord cultures leads to increases in ROS/RNS and c-Abl-P

To further analyze possible cellular mechanisms underlying MN death induced by mutSOD1-MCM, we focused on two classical pathogenic events reported in MNs in ALS models, namely accumulation of reactive oxygen species (ROS)/reactive nitrogen species (RNS) and activation of c-Abl (Fritz et al., 2013; Marchetto et al., 2008; Rojas et al., 2014, 2015). Based on our previous time-course imaging studies measuring ROS/RNS and c-Abl-P in spinal cord neurons induced by mutSOD1 astrocyte-conditioned media (Rojas et al., 2014, 2015), in the current study we applied mutSOD1-MCM for 90 min to 4 DIV spinal neurons for both sets of experiments. NTg-MCM and MN media were used as negative controls (90 min), and H_2_O_2_ (200 µM, 20 min) was used as positive control. To investigate whether mutSOD1-MCM leads to increases in intracellular ROS/RNS levels, following incubation, spinal cord neurons were washed and loaded for 30 min with CM-H_2_DCF-DA. This non-fluorescent dye passively diffuses into cells, but upon hydrolysis, the generated DCFH carboxylate anion is trapped inside the cells where oxidation leads to the formation of the fluorescent product DCF. The increased DCF fluorescent intensity reflects the accumulation of certain ROS/RNS species. Using combined real-time fluorescence and phase-contrast imaging, a substantial increase in intracellular DCF fluorescence was observed upon application of MCM-mutSOD1 (diluted 1/4 and 1/8) in spinal cord neurons, including in MN-like cells (Fig. 3A-B). The application of H_2_0_2_ mimicked this increase in DCF labeling. In control conditions, NTg-MCM and MN media did not change intracellular DCF levels (Fig. 3A-B).

**Figure 3.**
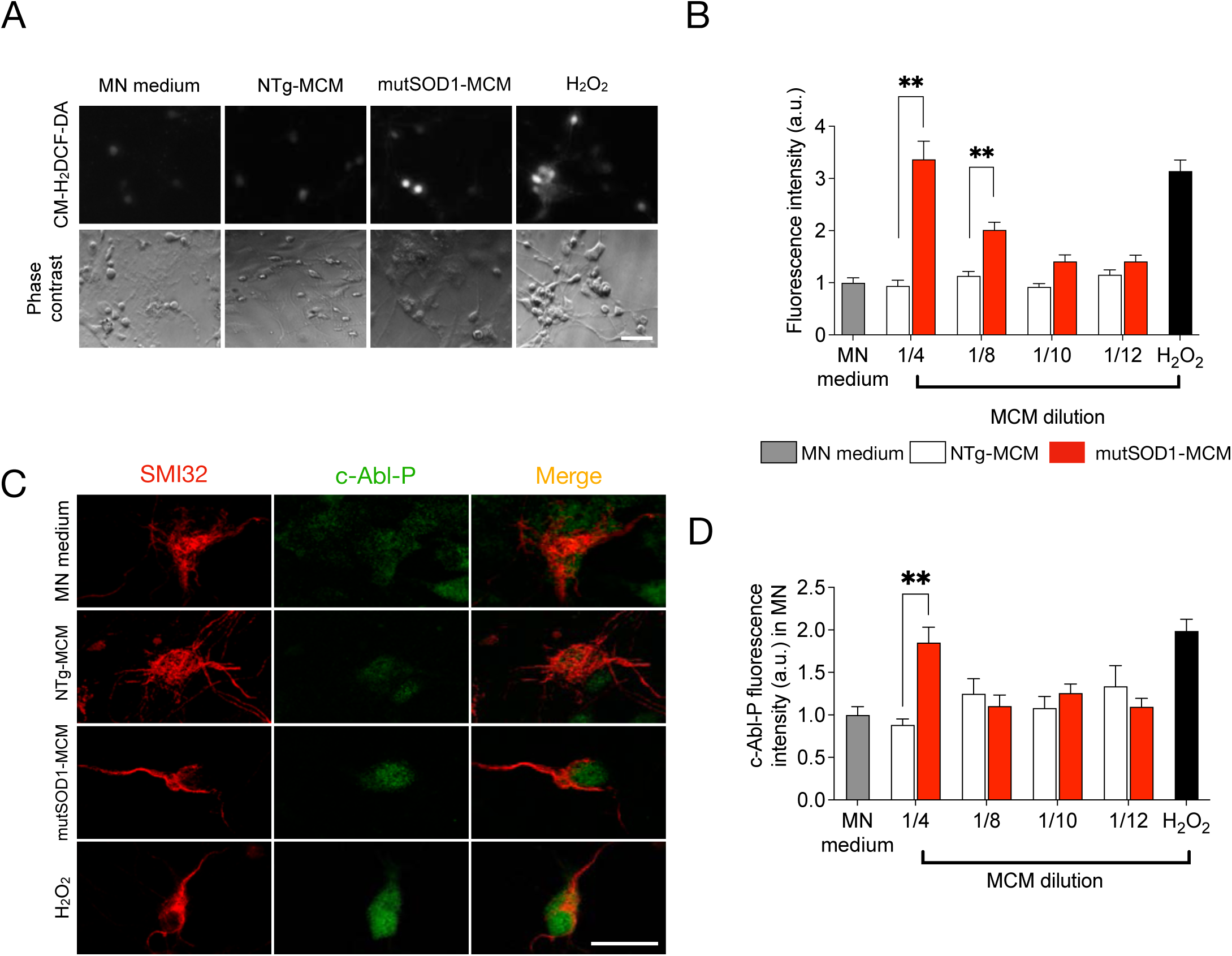
MCM-mutSOD1 triggers phosphorylated c-Abl and H_2_O_2_ accumulation. **A**, Representative images of DFC assay and phase contrast of NTg spinal cord cultures exposed to MCM-mutSOD1 (dilution 1/4), NTg-MCM (dilution 1/4), MN medium, and H_2_O_2_ (200 mM) as a positive control. Scale bar: 50 μm. **B**, Graph of the average intensity of DFC probe in neurons treated for 90 min with MCMs at different dilutions, as indicated. The graph represents the average ± s.e.m of 3 experiments performed independently and analyzed by one-way ANOVA (**, P <0.005 relative to NTg-MCM. **C**, Representative images of immunofluorescence against phosphorylated c-abl (c-Abl-P) and SMI32 (MNs) when exposed for 90 min to MCM-mutSOD1 (dilution 1/4), NTg-MCM (dilution 1/4), and MN medium, and H_2_O_2_ (200 mM, 20 min) as a positive control. Scale bar: 50 μm. **D**, Graphs showing fluorescence intensities (a.u.) for c-Abl-P at 4 DIV when NTg spinal cord cultures were treated acutely (90 min) with MCM at different dilutions, as indicated. The graph represents the mean ± s.e.m of 3 experiments performed independently and analyzed by one-way ANOVA (** P <0.005) relative to the NTg-MCM.

Using immunolabeling, we next evaluated the effect of mutSOD1-MCM on the levels of phosphorylated c-Abl kinase (c-Abl-P), a tyrosine kinase widely associated with neuronal apoptosis, activated under a wide range of stimuli including inflammation, DNA damage, amyloid beta, and oxidative stress (Etten, 1999; Klein et al., 2010; Martinez et al., 2023; Schlatterer et al., 2011; Tsai & Yuan, 2003; J. Y. J. Wang, 2005; Yáñez et al., 2016). After 90 min of application of the media, cells were fixed and double immunostained with an antibody against SMI32 to identify MNs and with a specific antibody that recognizes c-Abl that is phosphorylated on tyrosine 412 (Tyr412), a site that enhances c-Abl catalytic activity (Hantschel & Superti-Furga, 2004). We detected a significant induction of c-Abl-P in MNs incubated with 1/4 dilution of MCM-mutSOD1 compared to controls NTg-MCM or MN media (Fig. 3C-D). Together, these results indicate that soluble toxic factor(s) released by myotubes that carry mutSOD1 lead to increases in ROS/RNS and c-Abl-P in MNs.

### 3.4 MutSOD1-MCM induces MN death distally through the axons

Muscle-motoneuron communication is key for maintaining the NMJ and for the long-term survival of MNs (Ionescu et al., 2016). Alterations in intracellular communications can lead to synapse disruption and axon degeneration, which could be an inflection point in neurodegenerative diseases, including ALS (Maimon et al., 2018). The molecular signaling for the correct maintenance of neuromuscular communication can act locally at the synapse or travel long distances through the axon by retrograde transport. In our previous survival experiment (see Fig. 2), the MN soma and the axon are in direct contact with the MCM. Therefore, it is not possible to determine if the observed effects of myotube-derived toxic factor(s) on MN survival exert their impact in the soma (proximal effect) or through the axons (distal effect). For this reason, we used microfluidic devices that enable a physical separation between the neuronal soma and its synaptic terminal, with no fluid exchange between the chambers. Primary MNs from neonatal wild-type mice were cultured in the microfluidic devices in the proximal compartment (Fig. 4A, B). At 4 DIV, MCM 1/4 dilution was applied in the distal compartment (Fig. 4B). After three days, cells were fixed and immunostained for SMI32 and MAP2 to determine MN survival (Fig. 4C). To evaluate the survival of MNs more precisely, the MN compartment was divided into two sides (sides A and B; Fig. 4A), separating the MNs that were further from the microchannels, which were less likely to have direct contact with the MCMs (termed non-innervating side A), from those MNs closer to the microchannels and hence more likely to have innervated the distal chamber and thus directly contact the conditioned environment (termed innervating side B) (Fig. 4C). We found that distal application of mutSOD1-MCM reduced MN survival by 30% in the cell located in innervating side B, while no significant MN loss was detected in non-innervating side A (Fig. 4C-E). Moreover, we found that neither NTg-MCM nor MN media reduced MN survival on either side A or B (not shown). Together, these results indicate that the toxic factor(s) present in the mutSOD1-MCM exerts its effect retrogradely.

**Figure 4.**
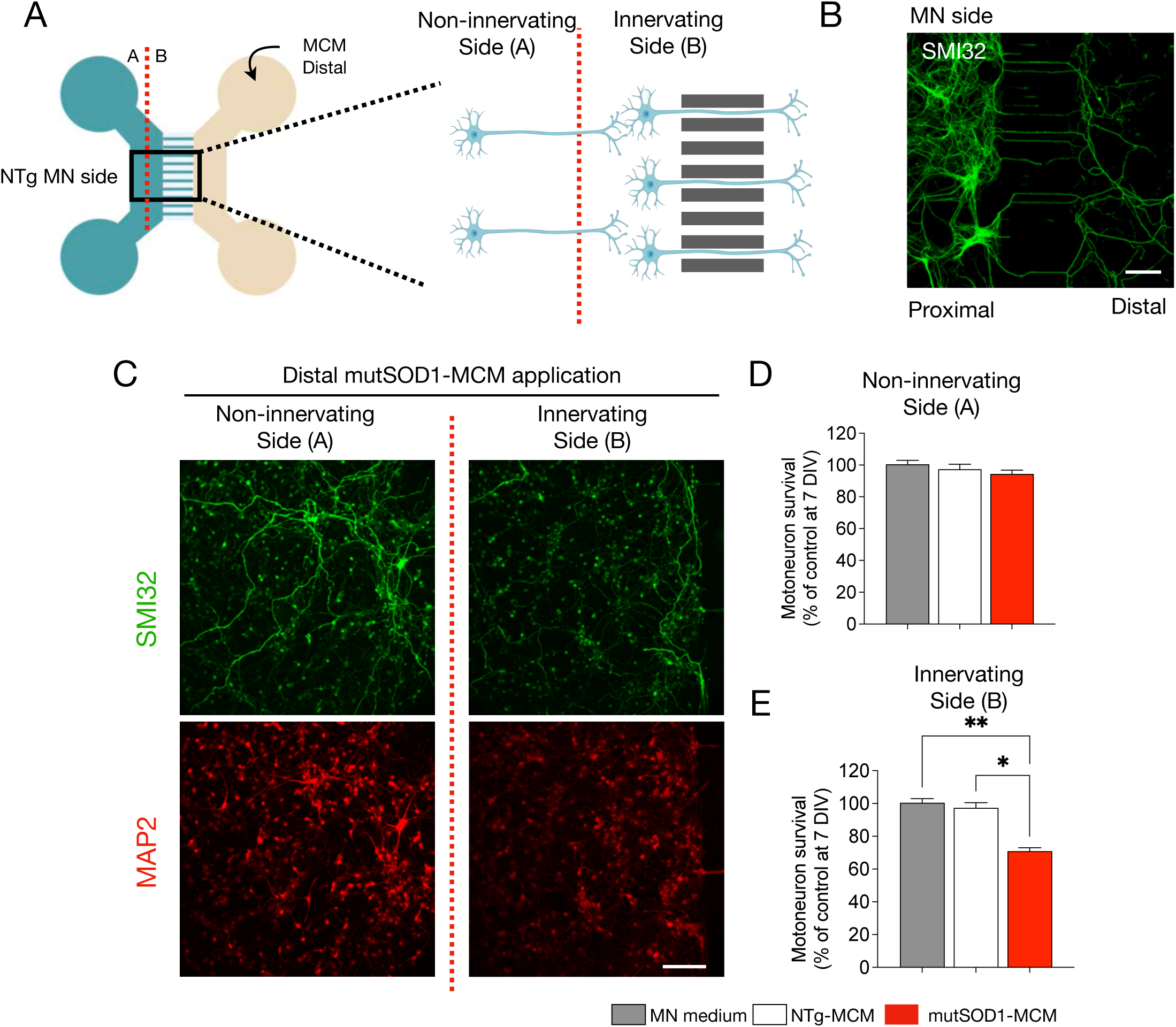
MN survival is reduced in microfluidic systems when exposed to mutSOD1-MCM. **A**, Representative diagram showing s microfluidic device to determine the survival of primary NTg MN cultures (cultured on MN side) at 14 DIV under the distal application of MCM-mutSOD1 or NTg-MCM for 3 DIV: indicated is the non-innervating (side A) and non-innervating (side B) of MNs. Next, MNs were fixed and incubated with specific antibodies against SMI32 to detect MNs and counted with respect to the total nuclei visualized with NucBlue staining. **B**, Immunostaining against SMI32 in 3 DIV NTg primary MN culture in a microfluidic device. Cells were plated in the proximal compartment and MCM-mutSOD1 was applied in the distal compartment for survival quantification. Scale bar: 200 µm. **C**, Representative images of immunofluorescence against SMI32 (MNs) and MAP2 (all neurons) when exposed to MCM-mutSOD1 in the non-innervating (side A) and non-innervating (side B) of the proximal chamber. Scale bar: 100 μm. **D**, Quantification of survival of non-innervating MN after distal treatment with MCM-mutSOD1 or NTg-MCM in a microfluidic chamber. Values represent the mean ± s.e.m. of 3 independent experiments and analyzed by one-way ANOVA relative to the NTg-MCM at 17 DIV. **E**, Graph of survival of innervating MN after distal treatment with MCM-mutSOD1 or NTg-MCM in the microfluidic chamber. Values represent the mean ± s.e.m. of 3 independent experiments and were analyzed by one-way ANOVA (*P <0.05 **P <0.005) relative to the NTg-MCM at 17 DIV relative to the control medium at 17 DIV.

### 3.5 Distal application of mutSOD1-MCM increases Ca^2+^ transients in wild-type MNs

Next, we wanted to get insights into molecular mechanisms that underlie the mutSOD1 myotube-mediated toxicity on MNs through a retrograde manner. To recreate the MN-muscle communication, and to ensure the generation of abundant functional MN axons in this distal chamber, WT (NTg) myotubes were grown in the distal compartment (Fig. 5A). Regarding potential mechanisms, we first focused on measuring intracellular Ca^2+^ transients as abnormalities in Ca^2+^ homeostasis has been implicated in the disruption of kinesin-mediated axonal trafficking (Hollenbeck & Saxton, 2005; Li et al., 2004). To determine if mutSOD1-MCM causes alterations in intracellular Ca^2+^ transients in the MNs, we used the GCaMP6 sensor to transduce the proximal MN compartment of the microfluidic devices (Rose et al., 2016). Seven days after transduction, we replaced the medium for MN media, NTg-MCM, or mutSOD1-MCM in the proximal and distal chambers (Fig. 5A). Five min later we recorded the MN somas for 2 min to analyze Ca^2+^ events. Independent of the chamber used, low MN activity (5-7 spontaneous Ca^2+^ events/min) was measured when MN media (control) or NTg-MCM was applied (Fig. 5B-C). By contrast, application of mutSOD1-MCM in the proximal chamber led to a significant increase of 2.5-fold (∼12 events/min) in transient Ca^2+^ events (Fig. 5B). When adding mutSOD1-MCM in the distal chamber, we observed an even higher increase of 3-4-fold (∼20 events/min) in transient Ca^2+^ events (Fig. 5C). These results show that soluble factor(s) released from mutSOD1 myotubes trigger an increase in Ca^2+^ transients, particularly when applied at the axonal compartment where the NMJ resides, suggesting a distinct spatial effect of toxic mutSOD1-MCM.

**Figure 5.**
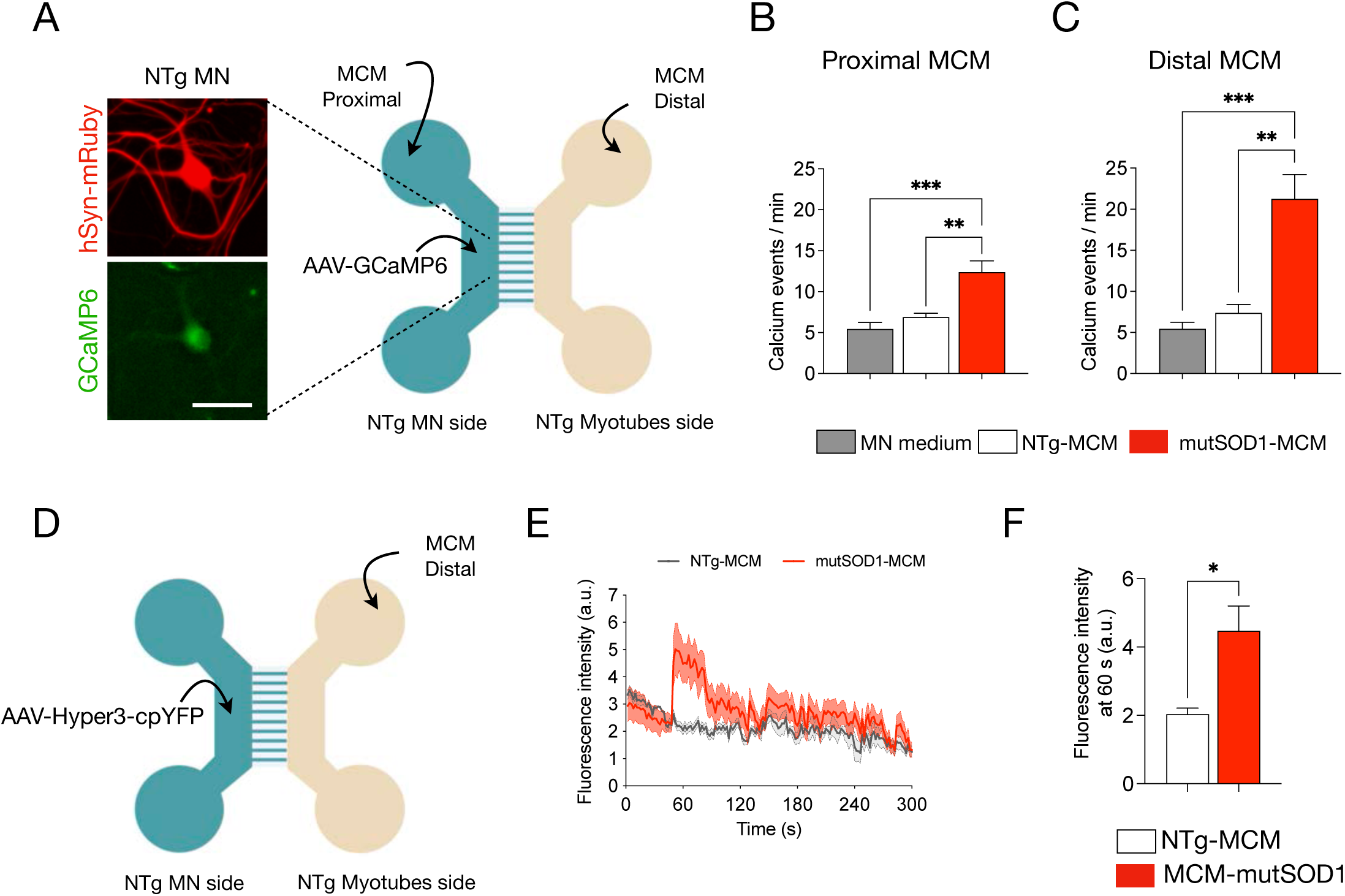
Application of mutSOD1-MCM rapidly increases calcium transients and induces H_2_O_2_ accumulation in wild-type MNs. **A**, Representative diagram of NTg MNs and NTg myotubes co-cultured in a microfluidic chamber. Images show examples of MN that was subjected to transduction with AAV1/2-hSyn-mRuby2-GCaMP6s (hSyn-mRuby in red, and GMaMP6s in green, left panel). Seven days later, cultures were exposed to NTg-MCM and MCM-mutSOD1 in the myotube (distal) or MN (proximal) compartment for 10 min before measuring calcium events. Scale bar: 50 µm. **B**, **C**, Quantification of the number of calcium events per min of MN exposed to the different MCMs, as indicated. Values of the graph represent the mean ± s.e.m. of 3 independent experiments and analyzed by one-way ANOVA (** P <0.005, *** P <0.0005) relative to the MN medium and NTg-MCM. **D**, Schematic of a co-culture of NTg MNs expressing the Hyper-3 sensor and NTg myotubes from P2 mice in a microfluidic chamber, where MCM (NTg and mutSOD1) were applied in the distal myotube compartment. **E**, Signal profile plot of Hyper3 fluorescent signal vs. time after application NTg-MCM and MCM-mutSOD1.Values of the graph represent the experimental average of 6 cells (n=6), using the same microfluidic chamber, and analyzed by t-student test (*P <0.05) relative to NTg-MCM. **F**, Fluorescent intensity of Hyper measured at 60 s from the profile, after application NTg-MCM and MCM-mutSOD1. Values represent the mean ± s.e.m. and analyzed by t-student test (*P <0.05) relative to NTg-MCM (n=6).

### 3.6 mutSOD1-MCM applied distally induces H_2_O_2_ accumulation in wild-type MNs

Studies in ALS-related SOD1 mutations indicate that mitochondrial dysfunction participates in the pathogenesis of MNs through the generation of intracellular ROS (Barber & Shaw, 2010; Rojas et al., 2014, 2015; Vehviläinen et al., 2014). As reported with DCF (Fig. 3), mutSOD1-MCM triggers an accumulation of ROS/RNS in WT (NTg) spinal cord cultures. To evaluate whether mutSOD1-MCM can induce ROS accumulation in MNs in a distal and retrograde manner in the microfluidic cultures, we used the HyPer-3 probe as a biosensor for intracellular H_2_O_2_ in living cells (Bilan et al., 2013). Co-cultures of wild-type myotubes and MNs were generated, with the latter neuronal compartment transduced with AAV1/2-HyPer-3. Seven days after infection, NTg-MCM or mutSOD1-MCM was applied to the distal chamber (Fig. 5D-E), and serial images were taken in an epifluorescent microscope (every 2 sec for 6 min). We found that the application of mutSOD1-MCM significantly increased ∼2.5-fold the HyPer-3 fluorescent intensity relative to NTg-MCM (Fig. 5E-F). These results show that applying mutSOD1-MCM to the axonal compartment triggers the accumulation of intracellular H_2_O_2_ levels in MN somas.

### 3.7 MCM-mutSOD1 affects antegrade and retrograde mitochondrial axonal transporting in MNs

To analyze the possibility of an axonal trafficking dysfunction as a pathological cell event, we evaluated the antegrade (from the soma to the axonal terminal) and retrograde (from the axon terminal to the soma) mitochondrial transport in MN-myotube microfluidic co-cultures (Fig. 6A) to evaluate mitochondria trafficking by kymograph quantification (Fig. 6B) after proximal (Fig. 6C) or distal (Fig. 6D) application of MCM-NTg and MCM-mutSOD1. To visualize MN axons, 3 DIV MNs were infected with an AAV1/2 harboring a red fluorescent protein (RFP) fused to a mitochondrial targeting sequence Cox8 (Fig. 6A). mutSOD1-MCM or NTg-MCM was applied to the myotube (distal) or MN (proximal) compartments and 24 h later mitochondrial movement in the axons of the MN was recorded. We tracked the mitochondrial axonal movement through the microchannels of the microfluidic chamber, thereby only quantifying axons in contact with the distal compartment. The generated videos were converted to kymograph images representing the movement of a particle over time. The resulting slopes show mitochondrial velocity (Fig. 6C-D). Compared to NTg-MCM, we observed that incubation of mutSOD1-MCM in the proximal compartment (MNs) significantly decreased the mitochondrial velocity of both the antegrade (from 0.55 μm/s to 0.3 μm/s) and retrograde movement (from 0.4 μm/s to 0.3 µm/s) (Fig. 6C, E). Conversely, application of mutSOD1-MCM to the distal chamber only affected anterograde, but not retrograde transport (Fig. 6D, F). These results indicate that mutSOD1-MCM affects differentially axonal mitochondrial transport, with the antegrade transport being the most affected by the muscle-released toxic factor(s).

**Figure 6.**
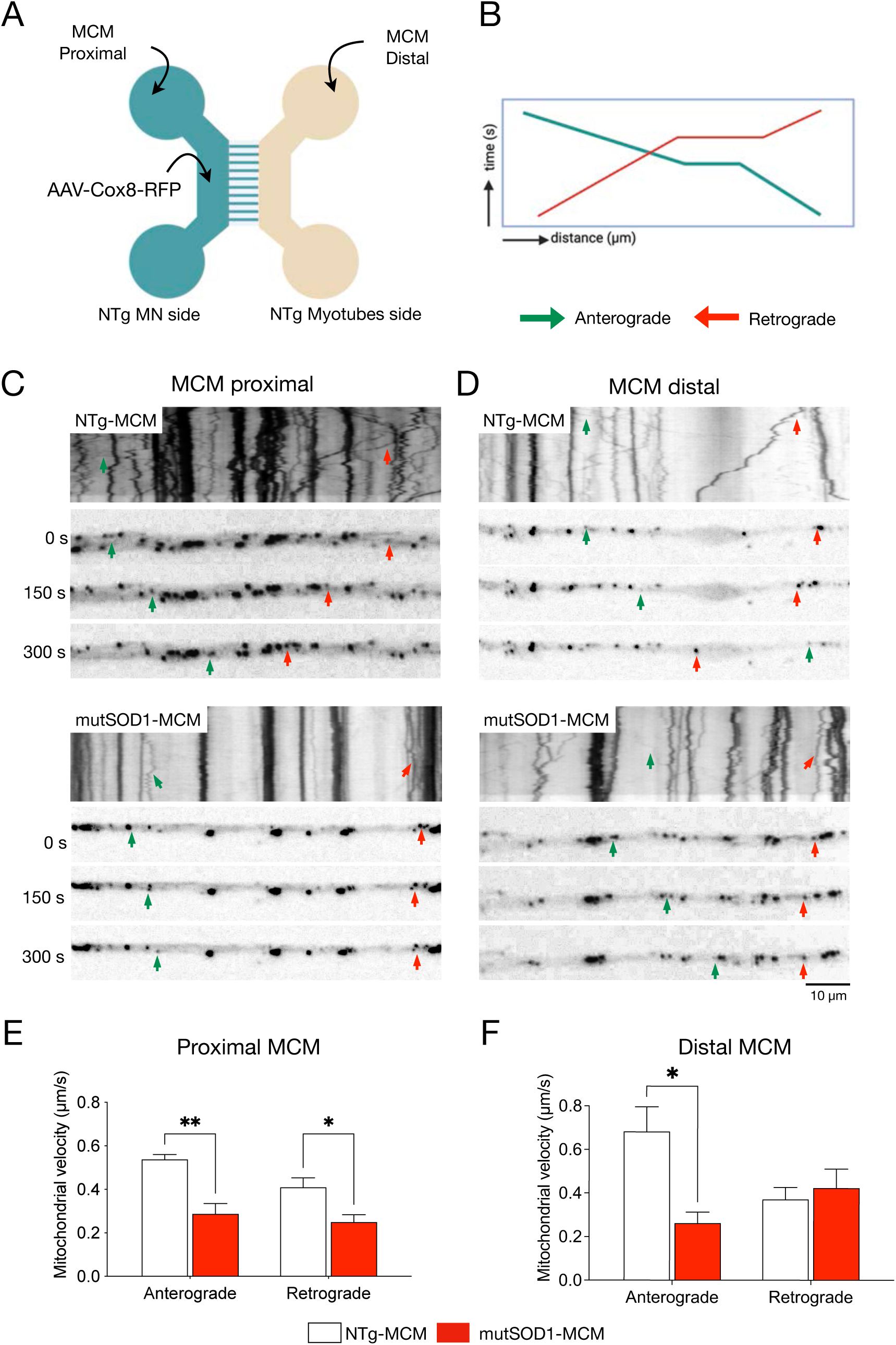
MCM-mutSOD1 applied proximally affects both antegrade and retrograde axonal transport. **A**, Representative diagram of MN and myotube co-cultures in a microfluidic chamber. MNs were subjected to transduction with AAV1/2-Cox8-RFP. Seven days later, cultures were exposed to NTg-MCM and MCM-mutSOD1 in the MN (proximal) or myotube (distal) compartment for 24 h for mitochondrial recording. **B**, Schematic of a kymograph to quantify mitochondria movements. Green lines and arrows indicate anterograde movements (from MN soma to axons), while red lines and arrows indicate retrograde movements (from axons to MN soma). **C**, **D**, Representative kymographs of mitochondrial movement in MN axons exposed to the different MCM, as indicated. **E**, **F**, Graphs of axonal mitochondrial velocity quantified from generated kymographs when MCM was applied in the proximal **(E**) and distal (**F**) chambers. Values of the graph represent the mean ± s.e.m. of 3 independent experiments and analyzed by t-Student test (*P <0.1, **P <0.01) relative to NTg-MCM.

## 4 DISCUSSION

The role of muscle in ALS pathology involves complex interactions, including potential contributions to disease progression and the modulation of MN function. Even though the role of skeletal muscle in ALS pathogenesis remains controversial, increasing evidence points out a preponderant non-autonomous mechanism underlying muscle degeneration (Badu-Mensah et al., 2020; Dobrowolny et al., 2005; Dupuis et al., 2009; Wong & Martin, 2010). To elucidate novel non-autonomous mechanisms of muscle-mediated MN degeneration in ALS, we generated a highly homogenous population of primary skeletal myotubes from myoblasts derived from ALS transgenic mice expressing human mutSOD1 and control NTg littermates. Characterization of the cultures revealed that mutSOD1 skeletal myotubes display phenotypic and functional differences compared to control myotube cultures. Given that our *in vitro* ALS muscle model is free of other critical implicated cell types, such as MNs, astrocytes, and microglia, our data suggest that the aberrant phenotypic and functional signature of mutSOD1 myotubes is cell autonomous. We also studied how conditioned media derived from mutSOD1 myotubes affects the function and survival of healthy NTg rodent MNs in typical mixed spinal cord cultures and compartmentalized microfluidic chambers enriched in MNs. Our finding that mutSOD1-MCM robustly kills MNs establishes that soluble toxic factor(s) released by ALS myotubes cause non-cell autonomous MN death. Furthermore, our study shows that mutSOD1-expressing myotubes exhibit phenotypic differences, suggesting a deaccelerated commitment from myoblasts to myotube differentiation. Specifically, we found that besides reduced myotube width, mutSOD1 myotubes display an increase in Pax7 and a decrease in MyoG protein expression. Our findings agree with a recent report where a human ALS skeletal muscle model was generated from induced pluripotent stem cells (iPSCs) derived from healthy individuals and ALS patients harboring mutations in SOD1 (Badu-Mensah et al., 2020). On the other hand, the expression pattern of Pax7 and MyoG is in stark contrast with data obtained from skeletal muscle biopsies of symptomatic ALS patients, where the expression of Pax7 was increased. In contrast, MyoG expression was reduced relative to control subject samples (Jensen et al., 2016). Based on additional data in the same study (comparing baseline with 12 weeks of progression), it was suggested that the activated myogenic process in symptomatic ALS muscle likely intends to overcome the denervation-induced muscle wasting. Comparing the data on cultured ALS myotubes with the analysis from skeletal muscle biopsies, we suggest that compensatory mechanisms during the progress of the disease are causing drastic alterations in myogenesis, changing from a deaccelerated process to an accelerated process. In future longitudinal studies, it would be interesting to determine underlying molecular mechanisms for the altered gene expression in pre-symptomatic and symptomatic ALS muscle cells.

The phenotypic alterations in cultured mouse and human mutSOD1 myotubes reproduce some of the validated muscle states *in vivo* models of ALS (Brooks et al., 2004; Hegedus et al., 2007, 2009; Manzano et al., 2011, 2013; Scaramozza et al., 2014; Wong & Martin, 2010). For instance, we further investigated the contraction frequency of our primary myotubes as a marker of contractile function. While several studies have reported impaired contractile function in mutSOD1 models, most employed electrical stimulation (not measuring spontaneous activity) and focused on adult mouse muscle or primary myotubes co-cultured with MNs (Badu-Mensah et al., 2020; Benlefki et al., 2020; Derave et al., 2003; Dupuis et al., 2004; Wier et al., 2019). Notably, Derave et al. (2003) observed slowed contraction in aged mutSOD1 mice at later stages of disease progression (P90 and P120). In contrast, our study uniquely assessed spontaneous contraction frequency in isolated myotubes derived from neonatal mice. Surprisingly, we found increased spontaneous contraction frequency in mutSOD1 myotubes compared to controls. This finding is intriguing, considering the delayed maturation observed in our mutSOD1 myotubes. It suggests that altered contractile behavior may manifest early in disease progression, independent of MN influence, and warrants further investigation into the underlying mechanisms.

Interestingly, we found that applying mutSOD1-MCM to the cultures further revealed a rapid induction of several classical pathogenic events in MNs, including impaired mitochondrial transport, disturbed calcium homeostasis, oxidative stress accumulation, and cell death induction. The findings underlying the mutSOD1-MCM application support previous observations of MN cell pathology which include impaired axonal transport of mitochondria from muscle to motor neurons contributing to ALS pathogenesis. Other studies demonstrate disrupted mitochondrial transport, calcium overload, and oxidative stress in ALS mouse models, leading to MN degeneration (Magrané et al., 2012; Rojas et al., 2014, 2015). Furthermore, dysregulation of calcium homeostasis in ALS involves non-cell-autonomous processes, as previously demonstrated by our lab using mutSOD1-ACM (Fritz et al., 2013). The observation of these altered processes mediated by toxic factor(s) present in mutSOD1-MCM may suggest a novel non-autonomous mechanism in ALS.

Soluble factor(s) released by mutSOD1 myotubes exert their toxic effects predominantly retrogradely, causing axonopathy and leading to lethal pathogenic changes. Our present *in vitro* evidence showing non-cell-autonomous toxic actions of mutSOD1 myotubes to healthy MNs agrees with several studies *in vitro* and *in vivo*. Thus, Maimon and colleagues (2018), using a compartmentalized microfluidic co-culture system with wild-type MN explants and primary myocytes, demonstrated that diverse ALS-causing genes, including mutations in SOD1, TDP43, and C9ORF72, promoted axon degeneration. In addition, studies using transgenic mice that express mutSOD1 selectively in skeletal muscles found alterations associated with ALS pathogenesis (Dobrowolny et al., 2009; Maimon et al., 2018; Wong & Martin, 2010). Specifically, Dobrowolny and colleagues (2009) showed that muscle-specific expression of mutSOD1 in mice induces severe muscle atrophy accompanied by microglia activation in the spinal cord but without evident signs of MN degeneration. Wong and Martin (2010) found that their transgenic mice exhibiting a skeletal muscle-restricted expression of mutSOD1 also developed muscle pathology and neurologic and pathologic phenotypes consistent with ALS, evidenced by spinal MNs developing distal axonopathy and significant MN degeneration. As indicated in the latter work, the difference between the severity of alterations in the spinal cord observed with the two transgenic studies could be explained by the aging of the animals; thus, Wong and Martin (2010) led their mice to become old, while Dobrowolny et al. (2009) performed their analyses on much younger mice (1–4-month-old) (Dobrowolny et al., 2009; Maimon et al., 2018; Wong & Martin, 2010). Despite these studies that support our findings, other *in vitro* and *in vivo* studies did not find evidence for the primary role of muscles in ALS. Specifically, in another *in vitro* study, conditioned media generated by myotubes/myocytes derived from the same transgenic mutSOD1 mouse model as we used to be unable to reduce the survival of NTg mouse MNs, either analyzed in mixed spinal cord cultures or cultures enriched for embryonic stem-cell derived MNs (Nagai et al., 2007). While the media was conditioned for seven days in both studies, the reason(s) underlying the difference between their results and ours may be related to technical issues associated with the generation of myotubes. For example, in our study, mutSOD1 myoblasts were differentiated into myotubes in 10 days, a process validated by phenotypic and functional analysis. In the previous study (Nagai et al., 2007), it was reported that myotubes were formed in 2-3 days from myoblasts (without showing characterization), making it plausible that not fully differentiated myotubes were generated from mutSOD1 myoblasts. Two *in vivo* studies also indicate that muscle is not a primary target for non-cell-autonomous toxicity in ALS (T. M. Miller et al., 2006; Towne et al., 2008). It was shown that delivery of RNA interferences (RNAi) targeting SOD1 to skeletal muscles in the mutant SOD1^G93A^ mouse model did not alter the time of onset of the disease or its progression despite causing a 50-60% reduction in SOD1 protein levels in the examined muscle (K. E. Miller & Sheetz, 2004; Towne et al., 2008). In both studies, the viral particles (AAV and lentivirus) to deliver RNAi against SOD1 were injected in young adult mutSOD1 mice; thus, intra-muscularly at P40 (T. M. Miller et al., 2006) or intravenously at P42 (Towne et al., 2008). Given that systematic studies of hindlimb muscles in mutSOD1 mice revealed functional and structural motor unit loss starting already at P40-P50 (Fischer et al., 2004; Frey et al., 2000; Hegedus et al., 2007; Saxena & Caroni, 2011; Zundert et al., 2012), it is plausible that the late viral delivery of SOD1-RNAi to the skeletal muscles was unable to significant protect and/or revert already induced muscle damage. This would not be surprising as multiple studies designed to reduce mutSOD1 in the CNS revealed that silencing in SOD1 gene expression only was able to significantly delay ALS onset and/or extend lifespan when the treatment was started during early developmental, strongly declining efficacy in maturing mice (Zundert & Brown, 2017).

Together, our data presented here, along with previous *in vitro* and *in vivo* studies (Badu-Mensah et al., 2020; Dobrowolny et al., 2009; Maimon et al., 2018; Wong & Martin, 2010), demonstrate that ALS skeletal muscle causes MN death and classic pathogenicity through non-cell-autonomous processes. Identifying the factors released by ALS skeletal muscle that are toxic to MNs will be essential to translate this knowledge into muscle-targeting treatments for ALS.

## DECLARATIONS

### Ethics approval

All experiments conducted in mice were handled according to the guidelines for the handling and care of experimentation established by the NIH (NIH, Maryland, USA) and following the protocol approved by the bioethics committee of Andres Bello University (approval certificate 014/2017).

### Availability of data and material

All data supporting our findings are in the manuscript. Any materials are available upon requesst.

### Authors’ contributions

P.M., F.J.B., and B.v.Z. conceived, designed the project, and wrote the manuscript. P.M., F.J.B., M.S., and S.A. performed experimental work. M.F.T., E.J., and M.C.-P. contributed to experimental design, manuscript review, and editing.

